# *Medicago truncatula Yellow Stripe-Like7* encodes a peptide transporter required for symbiotic nitrogen fixation

**DOI:** 10.1101/2020.03.26.009159

**Authors:** Rosario Castro-Rodríguez, María Reguera, Viviana Escudero, Patricia Gil-Díez, Julia Quintana, Rosa Isabel Prieto, Rakesh K. Kumar, Ella Brear, Louis Grillet, Jiangqi Wen, Kirunkamar S. Mysore, Elsbeth L. Walker, Penelope M. C. Smith, Juan Imperial, Manuel González-Guerrero

## Abstract

Yellow Stripe-Like (YSL) proteins are a family of plant transporters typically involved in transition metal homeostasis. The substrate of three of the four YSL clades (clades I, II, and IV) are metal complexes with non-proteinogenic amino acid nicotianamine or its derivatives. No such transport capabilities have been shown for any member of the remaining clade (clade III), which is able to translocate short peptides across the membranes instead. The connection between clade III YSL members and metal homeostasis might have been masked by the functional redundancy characteristic of this family. This might have been circumvented in legumes through neofunctionalization of YSLs to ensure a steady supply of transition metals for symbiotic nitrogen fixation in root nodules. To test this possibility, *Medicago truncatula* clade III transporter MtYSL7 has been studied both when the plant was fertilized with ammonium nitrate or when nitrogen had to be provided by endosymbiotic rhizobia within the root nodules. MtYSL7 is a plasma membrane protein expressed in the vasculature and in the nodule cortex. This protein is able to transport short peptides into the cytosol, although none with known metal homeostasis roles. Reducing *MtYSL7* expression levels resulted in diminished nitrogen fixation rates. In addition, nodules of mutant lines lacking *YSL7* accumulated more copper and iron, the later the likely result of increased expression in roots of iron uptake and delivery genes. The available data is indicative of a role of MtYSL7, and likely other clade III YSLs, in transition metal homeostasis.

**ONE SENTENCE SUMMARY:** *Medicago truncatula* YSL7 is a peptide transporter required for symbiotic nitrogen fixation in legume nodules, likely controlling transition metal allocation to these organs.

## INTRODUCTION

Iron, copper, zinc, and other transition metals are essential nutrients for plants (Marschner, 2011). These elements are structural components or cofactors of proteins involved in almost every physiological process. Therefore, obtaining transition metal nutrients from soil and delivering them to hundreds of metalloproteins in different organelles and tissues is essential for plant development. Several metal transport families participate in transition metal allocation (Pilon, 2011; Kobayashi and Nishizawa, 2012; Olsen and Palmgren, 2014). While most of them are evolutionary conserved in all domains of life, a few, such as the Yellow Stripe-Like (YSL) proteins, are exclusively present in plants (Curie et al., 2008; Kumar et al., 2017), suggesting selective pressures for specialized metal transport systems. YSLs are evolutionarily related to the Oligopeptide Transporter (OPT) family, sharing the capability of transporting amino acid substrates into the cell cytosol (Lubkowitz, 2011). The amino acid substrates of the biochemically characterized YSLs are typically the non-proteinogenic amino acid nicotianamine or nicotianamine-derived molecules (mugineic acids) in complex with transition elements (Schaaf et al., 2004; Aoyama et al., 2009).

The founding member of the family is *Zea mays YS1*. Mutants of this gene presented interveinal chlorosis in leaves (yellow stripes), caused by a deficiency in iron uptake (Beadle, 1929; Curie et al., 2001). ZmYS1 and its orthologues like OsYSL15 are the main iron uptake system from soil in grasses, participating in what has been named as Strategy II (in contrast to the Strategy I in non-grasses) (Kobayashi and Nishizawa, 2012). Monocots secrete nicotianamine-derived phytosiderophores to the rhizosphere, where they sequester iron and other transition elements (Roberts et al., 2004; Schaaf et al., 2004; Nozoye et al., 2011). Subsequently, YSL proteins mediate the uptake of these complexes into the root epidermal cells (Curie et al., 2001). However, this is not the only role of YSLs. Both in monocots and dicots, YSLs facilitate the transport of metal-nicotianamine complexes as part of a more general role in long-distance metal transport and intracellular transport (DiDonato et al., 2004; Waters et al., 2006). For instance, *Arabidopsis thaliana* YSL1 and YSL3 are responsible for iron uptake from the vascular tissues, redistribution from senescent leaves, and delivery to the seed (Waters et al., 2006; Chu et al., 2010). More recently, it has been proposed that YSL proteins also participate in the systemic control of iron homeostasis in plants (Kumar et al., 2017).

Based on their amino acid sequence, YSL transporters can be classified into four different groups (Yordem et al., 2011). Group I includes ZmYS1, AtYSL1, and AtYSL3. Group II is integrated by a number of YSL proteins that have intracellular localization. Best characterized AtYSL4 and AtYSL6 are located in the vacuoles and/or in chloroplast membranes, and are involved in the mobilization of metal stores (Conte et al., 2013; Divol et al., 2013). YSL Group IV is formed only by monocot proteins, including metal transporting OsYSL8 (Aoyama et al., 2009). While members from these three groups have been linked to transition metal homeostasis, no physiological function has been attributed to YSLs belonging to Group III.

Group III YSLs AtYSL7 and AtYSL8 are able to transport peptides into the cell (Hofstetter et al., 2013). This is used by plant pathogen *Pseudomonas syringae* pv *syringae* to introduce in the plant cell the virulence factor SylA, a tripeptide involved in proteasome inhibition that dysregulates the plant immune response (Groll et al., 2008). However, it is unlikely that this is the physiological role of Group III YSLs. Considering that all the other three YSL groups are involved in transition metal homeostasis (Curie et al., 2001; Conte et al., 2013), we hypothesize that Group III members would also be participating in the same physiological process. However, no metal-related phenotype of any mutant of these family members have been reported in *Arabidopsis*, perhaps as a consequence of the frequent functional redundancy of the YSL family (Waters et al., 2006; Divol et al., 2013). As an alternative, we have considered the model legume *Medicago truncatula* due to its ability to carry out symbiotic nitrogen fixation, a process that heavily relies on metal transport (González-Guerrero et al., 2016). This symbiotic process required the neofunctionalization of many genes (De Mita et al., 2014), and could have caused the loss of some of the functional redundancy that characterizes the YSL family.

Transition elements are essential for symbiotic nitrogen fixation as cofactors of many of the enzymes participating in this process, including the enzyme central to the conversion of N_2_ into NH_4_^+^, nitrogenase (Brear et al., 2013; González-Guerrero et al., 2014). Considering the high expression levels of many of the genes encoding these metalloenzymes, a substantial part of the metal incorporated into the plant is directed to the root nodules, oftentimes eliciting metal deficiency responses (Terry et al., 1991). Consequently, it is expected that dedicated metal transport systems have been adapted from pre-existing ones to ensure proper metal supply to the newly developed organs. In those cases where functional redundancy might exist, it could imply that only one of the redundant genes acquire a novel role. Here, we take advantage of this possibility to study *M. truncatula* YSL7 (*Medtr3g063490*), a Group III YSL family member with high expression in nodules important for plant growth in general and nitrogen fixation in particular.

## RESULTS

### *MtYSL7* is expressed in roots and in nodules

The *M. truncatula* genome contains nine *YSL* genes (*MtYSL1-MtYSL9*). Sequence comparison of the encoded proteins with known YSL proteins from monocots and dicots, showed that *M. truncatula* YSLs are distributed in the same groups as any other dicot (Fig. 1A). Four of them (MtYSL1-4) belong to Group I, one (MtYSL6) is clustered in group II, and four (MtYSL5, 7, 8, and 9) are in Group III. MtYSL7 is very similar to AtYSL7 and GmYSL7, and likely resulted from a duplication event with MtYSL8 o MtYSL9. Over 89% identity is shared among MtYSL7, MtYSL8, and MtYSL9.

**Fig. 1.**
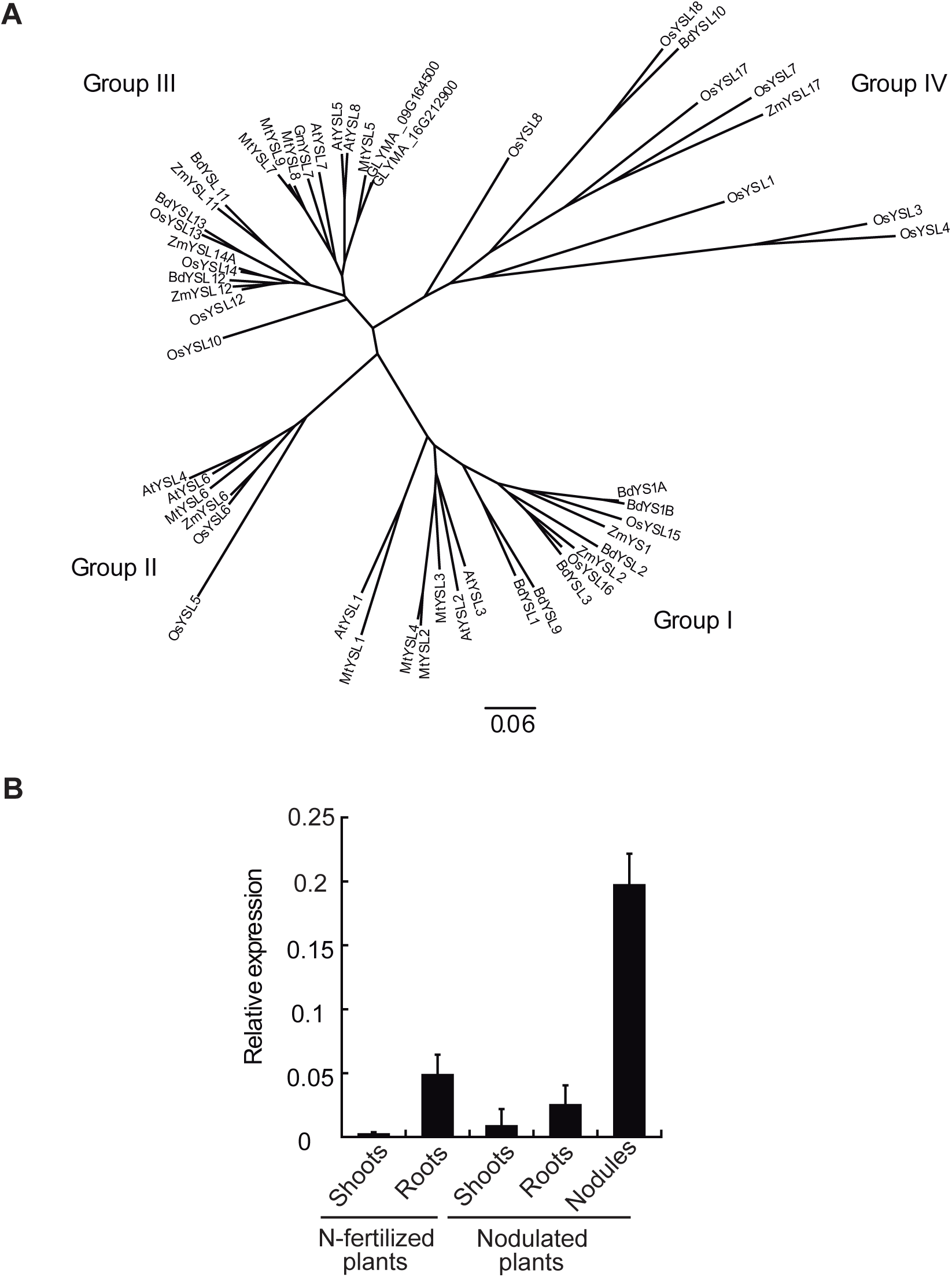
*Medicago truncatula YSL7* is a Group III YSL highly expressed in nodules. A) Unrooted tree of the *M. truncatula* YSL transporters, MtYSL1-MtYSL9 (*Medtr1g077840, Medtr1g007540, Medtr3g092090, Medtr1g007580, Medtr6g077870, Medtr7g028250, Medtr3g063490, Medtr5g01600*, and *Medtr3g063520*, respectively), and representative plant homologues. B) *MtYSL7* expression relative to internal standard gene *ubiquitin carboxyl-terminal hydrolase*. Data are the mean ± SE of five independent experiments.

*MtYSL7* was the only gene among the four members of the Group III *M. truncatula YSLs* to have a maximum of expression in nodules (Fig. 1B, Suppl. Fig. 1). Transcripts of these genes were specifically detected in roots and nodules, with no significant transcription in shoots. In spite of the high degree of similarity to *MtYSL7, MtYSL8* and *MtYSL9* were not expressed in nodules. In fact, no *MtYSL8* or *MtYSL9* transcripts were found in the organs at the time points analysed (Suppl. Fig. 1). In contrast, *MtYSL5* was expressed in all the organs tested, with the lowest relative expression being detected in nodules.

### MtYSL7 is a peptide transporter

YSL transporters have been typically associated with the transport of transition metal complexes with non-proteogenic amino acid nicotianamine or its derivates (Schaaf et al., 2004). To test whether MtYSL7 was able to transport transition elements free or complexed to nicotianamine, complementation assays of *S. cerevisiae* metal transport mutants were carried out. For this, the strains *ctr1* (deficient in copper uptake), *zrt1zrt2* (deficient in zinc uptake), and *fet3fet4* (deficient in iron uptake) were used (Askwith et al., 1994; Dancis et al., 1994; Dix et al., 1994; Zhao and Eide, 1996). These strains were co-transformed with a plasmid expressing a β-estradiol-dependent transactivator of *Gal* promoters, and a plasmid containing *MtYSL7* coding sequence regulated by a Gal promoter. Drop tests were carried out using serial dilutions of the cultures. In the case of iron, both the Fe^2+^ and the Fe^3+^ forms were tested. As shown in Suppl. Fig. 2, there was no restoration of the wild type growth in any of the conditions tested, unlike what has been reported for other metal-nicotianamine transporting YSLs in similar assays (Chu et al., 2010).

Alternatively, it can be hypothesized that the substrate of MtYSL7 is a peptide and not a metal-NA complex. YSL and OPT proteins are closely related (Lubkowitz, 2011), and the *Arabidopsis* AtYSL7 protein transports a peptide (Hofstetter et al., 2013). To test this possibility, different peptides were provided as the sole nitrogen source to an *S. cerevisiae* strain lacking an oligopeptide transporter (*opt1*) (Osawa et al., 2006) transformed with a plasmid containing *MtYSL7* (Fig. 2). *MtYSL7*-expressing yeasts were able to grow when four to six amino acid peptides were provided. Some limited growth was also observed when a 12 amino acid peptide was used as substrate, but not when one of 10 was provided. MtYSL7 did not appear to transport the tripeptide glutathione when expressed in the yeast glutathione transport mutant *hgt* (Suppl. Fig. 3). In addition, no evidence for transport of iron-regulatory peptide IMA (Grillet et al., 2018) was observed (Suppl. Fig. 4).

**Fig. 2.**
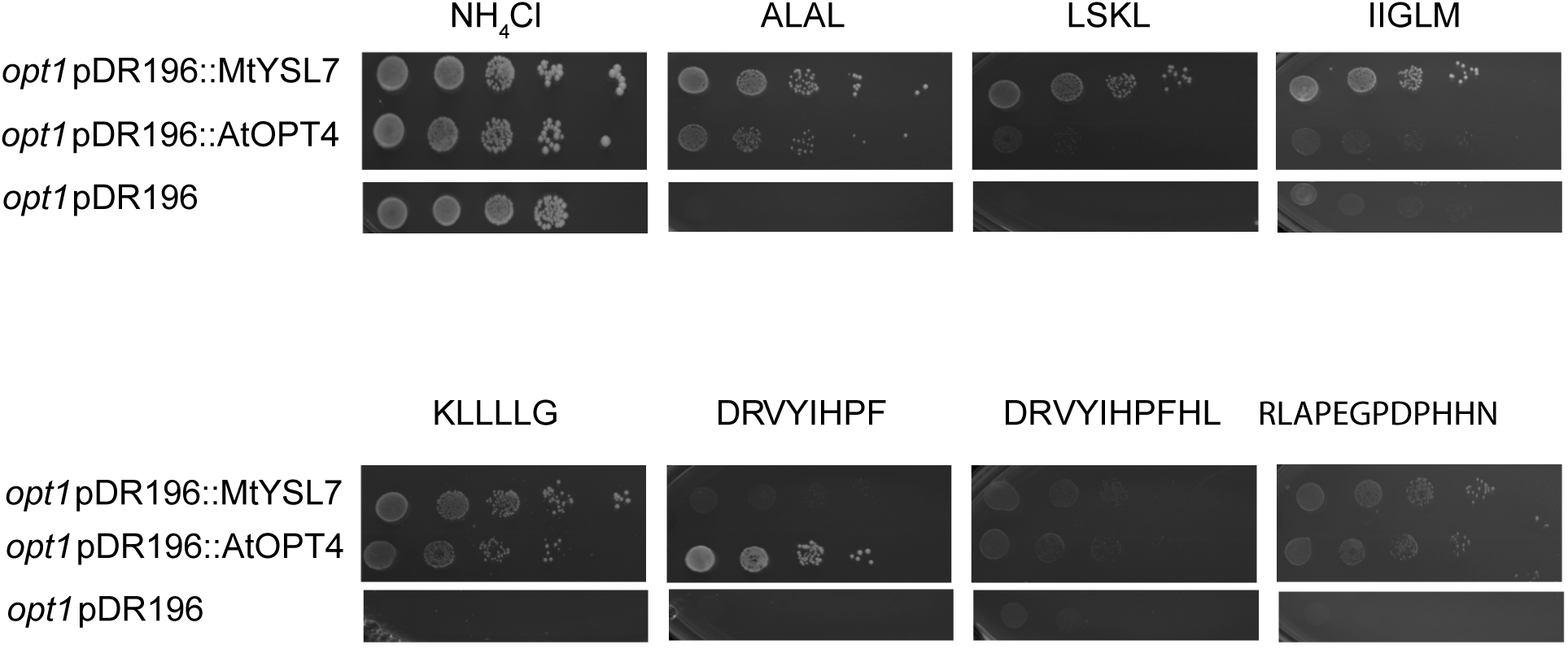
MtYSL7 transports peptides. The yeast strain *opt1*, mutant in the oligopeptide transporter ScOPT1, was transformed with either the empty pDR196 vector, or with pDR196 containing the coding sequence of *MtYSL7* or *AtOPT4*. Serial dilutions (10x) were grown on SD media supplemented with the nitrogen sources indicated.

### MtYSL7 is located in the plasma membrane of root pericycle and nodule cortical cells

To determine the tissue localization of the expression of *MtYSL7, M. truncatula* seedlings were transformed with a construct fusing the 2 kb region upstream of *MtYSL7* to the β-glucuronidase gene (*gus*). GUS activity was localized in 28 dpi plants based on the blue stain resulting from using X-Gluc as a substrate (Fig. 3). As indicated by the RT-PCR studies, *MtYSL7* was expressed in roots and nodules, with the signal being more intense in nodules (Fig. 3A). In nodules, *MtYSL7* expression was located in the cortical region of the nodule throughout its length (Fig. 3B, C), with no expression observed in the nodule core, regardless of the developmental zone of the nodule. In roots, GUS activity was detected in the perivascular region, in a zone that contains the endodermis or pericycle surrounding the vessels (Fig. 3D).

**Fig. 3.**
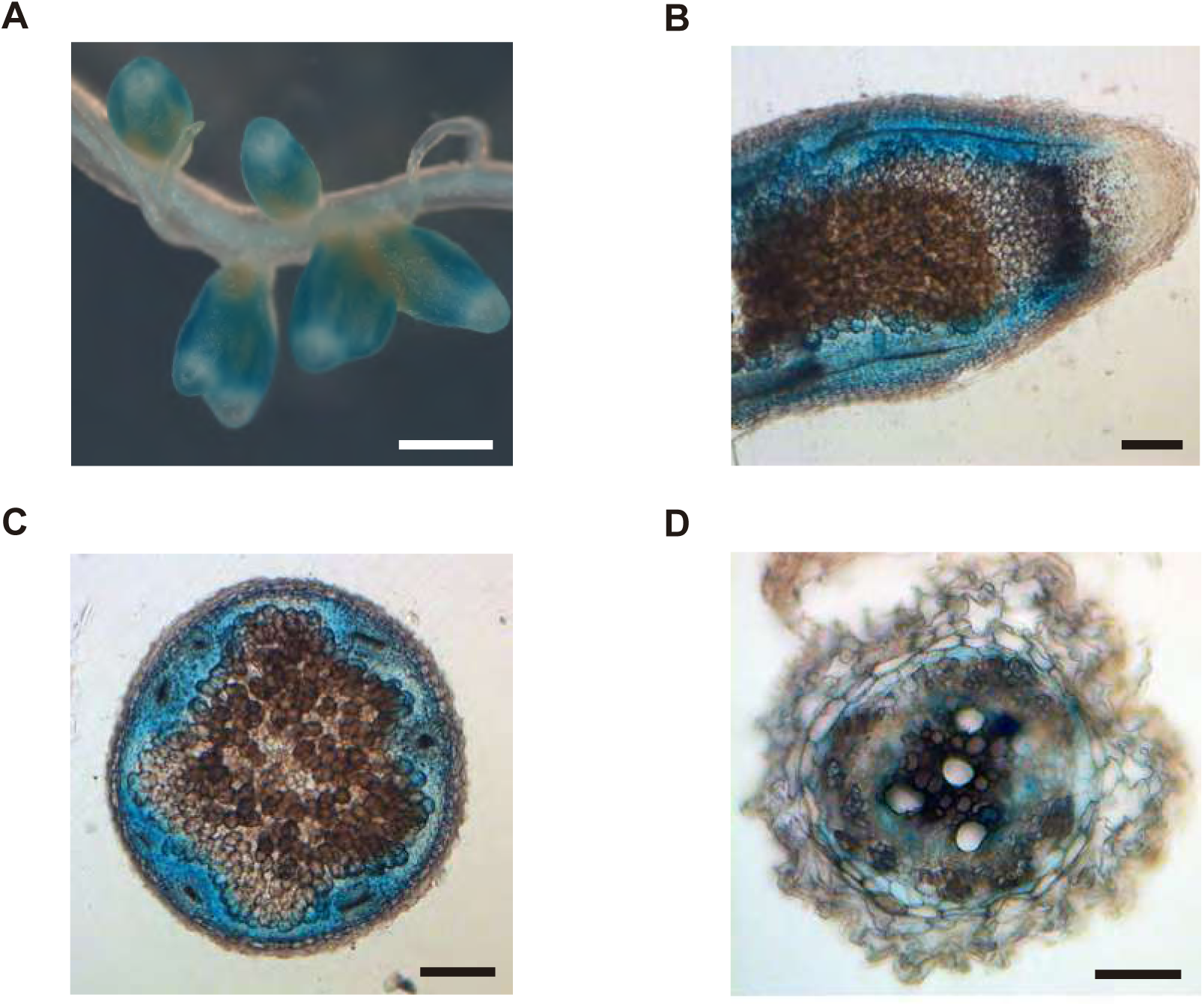
*MtYSL7* is expressed in the root and nodule vasculature and in the nodule cortex. A) Histochemical staining of the GUS activity in 28 dpi root and nodules expressing the *gus* gene under the regulation of the *MtYSL7* promoter. Bar = 1 mm. B) Longitudinal section of a GUS-stained 28 dpi nodule expressing *gus* under the *MtYSL7* promoter. Bar = 200 μm. C) Cross section of a GUS-stained 28 dpi nodule expressing *gus* under the *MtYSL7* promoter. Bar = 200 μm. D) Cross section of a GUS-stained 28 dpi root expressing *gus* under the *MtYSL7* promoter. Bar = 100 μm.

Immunolocalization of a C-terminal (HA)_3_-tagged MtYSL7 regulated by its own promoter supported the reporter gene studies (Fig. 4). The signal from the Alexa 594-conjugated antibody used to localize MtYSL7-HA was observed in the periphery of the nodule, in the cortical area, both in the perivascular and intervascular areas (Fig. 4A, B). At a higher magnification, the perivascular distribution of MtYSL7-HA seemed to be confined to the endodermal layer (Fig. 4C), mostly to the periphery of the cell. In roots MtYSL7-HA was observed in the pericycle (Fig. 4D). The intracellular distribution of MtYSL7-HA was indicative of a plasma membrane association, as also evidenced by its co-localization with a plasma membrane marker when co-transfected into tobacco leaves (Fig. 4E). Controls in which no Alexa 594-conjugated antibody was used showed no signal in the measured channels (Suppl. Fig. 5).

**Fig. 4.**
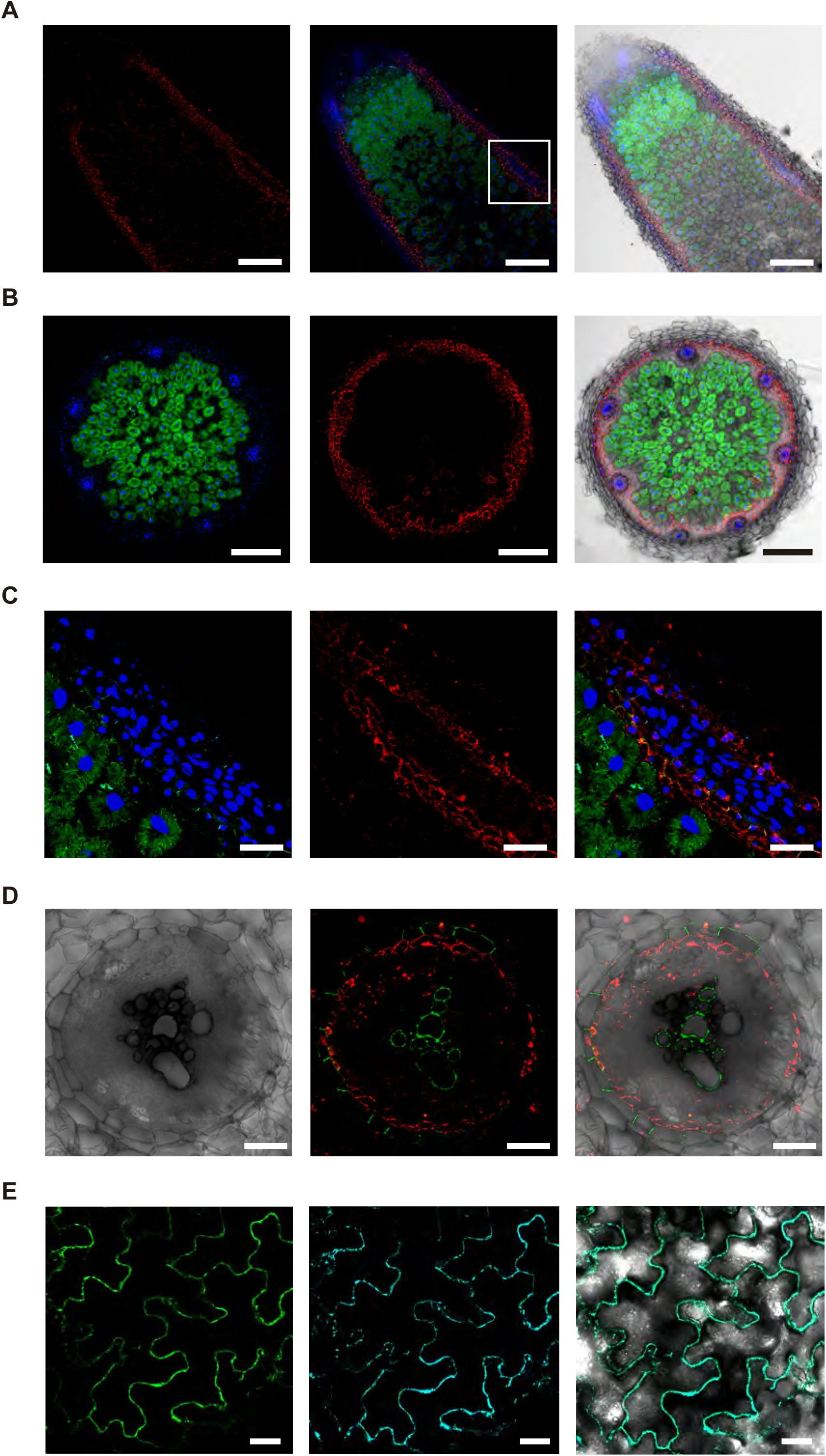
MtYSL7 is located in the periphery of nodule cortical cells, in the nodule endodermis and in the root pericycle. A) Longitudinal section of a 28 dpi *M. truncatula* nodule expressing *MtYSL7* under its own promoter and fused to three HA epitopes. The HA-tag was detected with the help of a secondary Alexa594-conjugated antibody (red). Transformed plants were inoculated with a GFP-expressing *S. meliloti* (green) and DNA was stained with DAPI (blue). Right panel shows the overlay of all these channels together with the bright field image. Bars = 200 μm. B) Cross section of a 28 dpi *M. truncatula* nodule expressing *MtYSL7* under its own promoter and fused to three HA epitopes. The HA-tag was detected with the help of a secondary Alexa594-conjugated antibody (red). Transformed plants were inoculated with a GFP-expressing *S. meliloti* (green) and DNA was stained with DAPI (blue). Right panel shows the overlay of all these channels together with the bright field image. Bars = 200 μm. C) Detail of a longitudinal section of a vessel from a 28 dpi *M. truncatula* nodule expressing *MtYSL7* under its own promoter and fused to three HA epitopes. The HA-tag was detected with the help of a secondary Alexa594-conjugated antibody (red). Transformed plants were inoculated with a GFP-expressing *S. meliloti* (green) and DNA was stained with DAPI (blue). Right panel shows the overlay of all these channels. The arrowheads indicate the position of the Casparian strip. Bars = 50 μm. D) Cross section of a 28 dpi root expressing *MtYSL7* under its own promoter and fused to three HA epitopes. The HA-tag was detected with the help of a secondary Alexa594-conjugated antibody (red). Lignin autofluorescence was used to identify xylem and Casparian strip lignin (green). Right panel shows the overlay of all these channels together with the bright field image. Bars = 50 μm. E) Colocalization of MtYSL7-GFP and AtPIP2-CFP in tobacco leaves. Left panel shows the localization of MtYSL7 fused to GFP (green) transiently expressed in tobacco leaf cells. Middle panel shows the localization of plasma membrane marker AtPIP2 fused to CFP, transiently expressed in the same cells. Right panel is the overlay of the two previous channels together with the bright field image. Bars = 50 μm.

### *MtYSL7* mutation affects plant growth and symbiotic nitrogen fixation

To determine the function of MtYSL7, two *M. truncatula Tnt1* insertional mutants were obtained from the Noble Research Institute (Tadege et al., 2008). NF11536 (*ysl7-1*) is a knock-down line with the transposon inserted in its first intron (nucleotide +1118) (Fig. 5A). The *Tnt1* insertion in NF9504 (*ysl7-2*) is in the first exon in nucleotide +315, what reduces *MtYSL7* expression below our detection limit. Regardless the extent to which expression is reduced, when uninoculated and grown with ammonium nitrate in the medium, both *MtYSL7* mutant alleles had reduced growth compared to wild type plants (Fig. 5B, C). The growth phenotype, particularly in roots, was restored in the knock-out *ysl7-2* mutant allele when a wild type copy of *MtYSL7* expressed under the control of its own promoter was reintroduced (Suppl. Fig. 6). In addition, although it was not evident from *in visu* examination of the plants, both mutant alleles had a slight but significant decrease in chlorophyll content (Fig 5D). In general, no significant differences in iron, copper, or zinc concentrations were observed, the one exception being copper levels in shoots of knock down *ysl7-1* (Fig. E-G).

**Fig. 5.**
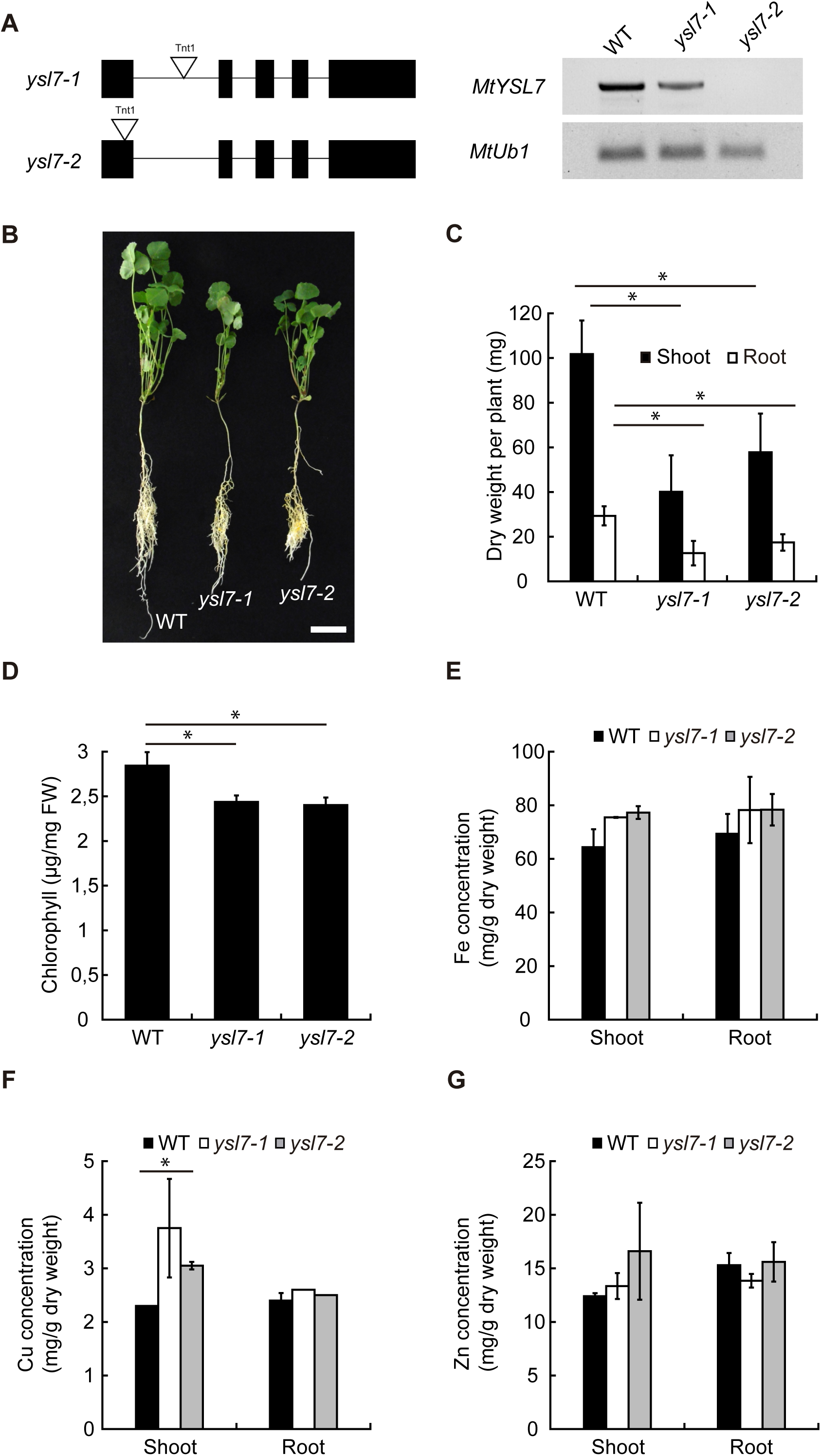
*MtYSL7* mutation affects plant growth under non-symbiotic conditions. A) Position of the *Tnt1* insertions in *ysl7-1* and *ysl7-2* lines. Lower half, RT-PCR of *MtYSL7* in 28 dpi nodules of wild type, *ysl7-1*, and *ysl7-2* lines. *Ubiquitin carboxyl-terminal hydrolase* (*MtUb1*) was used as a positive control. B). Growth of representative plants. Bar = 3 cm. C) Dry weight of shoots and roots of wild type, *ysl7-1*, and *ysl7-2* plants. Data are the mean ± SE (n = 5 plants). C) Chlorophyll content of wild type, *ysl7-1*, and *ysl7-2* shoots. Data are the mean ± SE of three sets of five pooled plants.E) Iron content in roots and shoots of wild type, *ysl7-1*, and *ysl7-2* plants. Data are the mean ± SE of three sets of five pooled plants. F) Copper content in roots and shoots of wild type, *ysl7-1*, and *ysl7-2* plants. Data are the mean ± SE of three sets of five pooled plants. G) Zinc content in roots and shoots of wild type, *ysl7-1*, and *ysl7-2* plants. Data are the mean ± SE of three sets of five pooled plants. * indicates statistical significance (p < 0.05).

Similar growth differences were observed when all the nitrogen was provided by symbiosis with *S. meliloti* (Fig. 6). Plant growth was reduced in *MtYSL7* mutants, at a higher degree in the knock-out line than the knock-down line (Fig. 6A, B). These plants developed nodules that had no evident morphological differences from wild type (Fig. 6C). Nitrogenase activity was affected in both *Tnt1* lines, with around 40% of the activity of controls (Fig. 6D). These phenotypes were restored in *ysl7-2* when a wild type copy of *MtYSL7* expressed by its native promoter was reintroduced (Suppl. Fig. 7). In addition, *ysl7-2* nodules had a significantly higher iron and copper content in nodules than wild type (Fig. 6E, F), while no differences were observed for zinc (Fig. 6 G). Increased iron level might be the result of the increased expression in roots of iron uptake and delivery systems, such as ferroreductase FRO1 and iron transporter MtNramp1 (Suppl. Fig. 8). Moreover, the increased iron content in *ysl7-2* nodules did not result in altered iron distribution in nodules (Supp. Fig. 9). Knock-down *ysl7-1* showed a similar, although non-significant metal accumulation trend, the likely consequence of the remaining *MtYSL7* expression (Fig 6E-G). In spite of the altered iron and copper homeostasis in nodules, reducing or increasing iron or copper content in the nutrient solution did not have any significant effect on plant growth or nitrogenase activity (Suppl. Fig. 10, Suppl. Fig. 11).

**Fig. 6.**
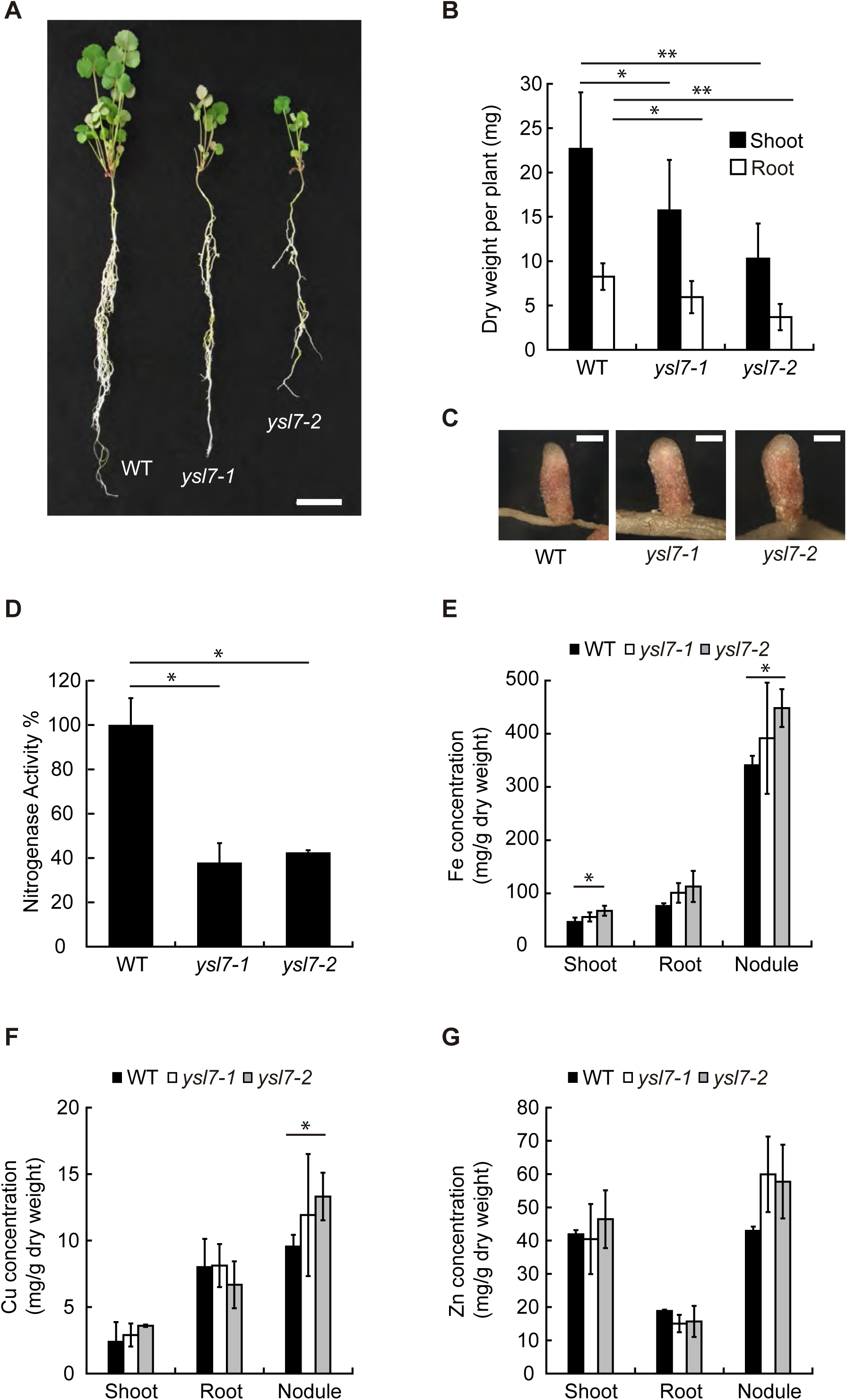
MtYSL7 participates in symbiotic nitrogen fixation. A) Growth of representative wild type, *ysl7-1*, and *ysl7-2* plants. Bar = 3 cm. B) Dry weight of shoots and roots of 28 dpi plants. Data are the mean ± SE (n = 10-15 plants). C) Detail of representative nodules of 28 dpi wild type, *ysl7-1*, and *ysl7-2* plants. Bars = 1 mm. D) Nitrogenase activity in 28 dpi nodules from wild type, *ysl7-1*, and *ysl7-2* plants. Data are the mean ± SE measured in duplicate from two sets of five pooled plants. 100 % = 0.28 nmol ethylene h^-1^ plant^-1^. E) Iron content in roots and shoots of wild type, *ysl7-1*, and *ysl7-2* plants. Data are the mean ± SE of two sets of five pooled plants. F) Copper content in roots and shoots of wild type, *ysl7-1*, and *ysl7-2* plants. Data are the mean ± SE of two sets of five pooled plants. G) Zinc content in roots and shoots of wild type, *ysl7-1*, and *ysl7-2* plants. Data are the mean ± SE of two sets of five pooled plants. * indicates statistical significance (p < 0.05).

## DISCUSSION

YSLs play a role in transition metal uptake from soil, its distribution from source to sink tissues, as well as in intracellular metal remobilization (Curie et al., 2001; Waters et al., 2006; Conte et al., 2013). More recently, there is evidence of YSL proteins participating in long-distance signalling of the nutritional status of iron (Kumar et al., 2017). These functions are carried out by members of three of the four known YSL groups (Yordem et al., 2011). The remaining one, Group III, has been associated with the transport of the *Pseudomonas syringae pv. syringae* virulence factor SylA (Hofstetter et al., 2013). Although it is unlikely that this is its physiological role, it showcases the ability of proteins from this family to use peptides instead of non-proteogenic amino acids as their substrate. Further reinforcing this role as peptide transporter, expression of *MtYSL7* in yeast restores the uptake capacity of short peptides. Similar transport capabilities were observed in GmYSL7 (accompanying manuscript). However, there seems to be some specificity to transport by MtYSL7, since a decapeptide was not transported, as neither were tripeptide glutathione or IMA-related peptides consisting of four, five, or seventeen amino acids.

*MtYSL7* is expressed in roots and nodules, with the highest levels in the nodules. Inmunolocalization of HA-tagged MtYSL7 indicates that it is associated to the plasma membrane of cells in the vasculature and in the nodule cortex and vasculature, similar to the subcellular localization of AtYSL7 (Hofstetter et al., 2013). In spite of being expressed in two different organs, it is in the nodules where MtYSL7 would be playing a primary role. In contrast to the mild reduction of growth and even milder lower chlorophyll content of *MtYSL7* mutants when plants do not need to develop nodules, reducing the expression levels of this gene leads to a diminished nitrogenase activity and a reduced plant growth in nodulating plants. This degree of neofunctionalization of MtYSL7 has allowed us to study the physiological role of Group III YSLs while avoiding possible functional redundancies common to this family (Waters et al., 2006; Divol et al., 2013). This trend is further developed in *G. max*, where orthologue GmYSL7 is specifically expressed in nodules and located in the symbiosome membrane (accompanying manuscript), suggesting a higher degree of specialization. However, GmYSL7 seems to be functionally equivalent to MtYSL7, since it is also able to complement the *ysl7-2* phenotype when its expression is driven by the *MtYSL7* promoter (accompanying manuscript). Based on *MtYSL7* expression in the nodule cortex, it could be hypothesized that MtYSL7 is introducing a yet-to-be determined metal complex into a cell layer with a specific need for metals and a role in preventing oxygen diffusion into the nodule. This would create a separate iron pool from those elements directed for nitrogen-fixing cells, which very likely is in the form of iron-citrate (Tejada-Jiménez et al., 2015; Kryvoruchko et al., 2018). However, if this were the case, we should expect that metal fortification of the nutrient solution would partially complement the phenotype, as reported for other nodule metal transport mutants (Tejada-Jiménez et al., 2015; Tejada-Jiménez et al., 2017; Gil-Díez et al., 2019). Moreover, we should also observe metal accumulation in the apoplast around these cells. The fact that this did not happen could be explained either as the absence of an additional low affinity metal uptake system in those cells or that MtYSL7 could be playing a differential physiological function.

An alternative hypothesis to the role of *MtYSL7* is that its substrate would serve as a signal of the metal status. In the case of soybean symbiosomes, it could reflect the metal condition of bacteroids within, while in *M. truncatula* nodule cortex, it would be an indicator of the available metal within the nodule. Supporting this theory is the observed increased iron and copper content of *ysl7-2* nodules, perhaps the consequence of lacking a feedback signal of metal sufficiency in nodules. Since no change in iron distribution was observed, it could be speculated that no defects in overall metal transport would result from *MtYSL7* loss-of-function. Consistent with this hypothesis is the upregulation of iron uptake systems in roots of *ysl7-2. MtFRO1* is a ferroreductase involved in the iron deficiency response in roots, responsible for the conversion of Fe^3+^ to Fe^2+^ prior to its assimilation (Andaluz et al., 2009). *MtFRO1* upregulation in roots should be the result of iron deficiency in the plant. However, we have not observed any significant reduction on iron concentration in the plant, but the contrary, with higher iron concentrations in *ysl7-2* nodules. Similarly, *MtNramp1* expression levels were higher in the roots of *ysl7-2* plants than in the controls, while no statistical differences were observed in nodules. This transporter is responsible for iron uptake by the root endodermis and by rhizobia-infected nodule cells (Tejada-Jiménez et al., 2015). The fact that it is upregulated exclusively in roots would be indicative of a signal controlling whole plant metal iron allocation instead of specifically being targeted to nitrogen-fixing cells. Furthermore, a more through transcriptomic approach in mutant orthologue *GmYSL7*, also indicate a large dysregulation of iron homeostasis (accompanying manuscript). Regardless of the specific cause of the induction of Fe deficiency responses in the root, these results link Group III YSL function to metal homeostasis, a physiological role shared by all the other three YSL groups (Curie et al., 2001; Aoyama et al., 2009; Conte et al., 2013), that could be suggested but had not been demonstrated previous to the research presented here. However, to conclusively prove any of these alternative theories, future work should focus on the identification of the specific substrate of Group III YSLs, for which a new set of metal-chelator peptides need to be identified.

## MATERIALS AND METHODS

### Plant growth conditions

*Medicago truncatula* R108 (Wild Type) and *Tnt1*-insertion mutants *ysl7-1* (NF11536) and *ysl7-2* (NF9504) seeds were scarified, sterilized and germinated as indicated by Tejada-Jiménez et al. (Tejada-Jiménez et al., 2015). Seedlings were planted on sterile perlite pots, and inoculated with *Sinorhizobium meliloti* 2011 or the same bacterial strain transformed pHC60-GFP (Cheng and Walker, 1998). Plants were grown in a greenhouse under 16 h light / 8 h dark at 25 °C / 20 °C conditions, and watered every two days alternating Jenner’s solution with water (Brito et al., 1994). Nodules were collected at 28 days-post-inoculation (dpi). Non-nodulated plants were grown under the same conditions of light and temperature but were watered every two weeks with solutions supplemented with 2 mM NH_4_NO_3_.

Hairy-root transformations of *M. truncatula* seedlings with *Agrobacterium rhizogenes* ARqua1 carrying the appropriate binary vector, were performed following the methodology described by (Boisson-Dernier et al., 2001).

### Yeast complementation assays

To analyse MtYSL7 peptide uptake capabilities, *MtYSL7* was amplified using the primers listed in Suppl. Table 1 and cloned in the *PstI* and *XhoI* sites of pDR196. Yeast strains YJL212C (OPT1) (Euroscarf) (BY4741; *MATa; ura3Δ0; leu2Δ0; his3Δ1; met15Δ0; YJL212c::kanMX4*) and BY4741 (Winston et al., 1995) were used. Growth experiments were performed using SD media without nitrogen (N) using the different peptides indicated as a nitrogen source. SD media supplemented with 5 g/l (NH_4_)_2_SO_4_ was used as a growth control media. Ten-fold serial dilutions were spotted (5 µl) onto SD or YPD plates and incubated at 28 °C for 2-3 days.

### Quantitative real-time RT-PCR

Gene expression studies were carried out by real-time RT-PCR (StepOne plus, Applied Biosystems) using the Power SyBR Green master mix (Applied Biosystems). Primers used are indicated in Suppl. Table 1. RNA levels were normalized by using the *ubiquitin carboxy-terminal hydrolase* gene as internal standard for *M. truncatula* genes (Kakar et al., 2008). RNA isolation and cDNA synthesis were carried out as previously described (Tejada-Jiménez et al., 2015).

### GUS staining

Two kilobases upstream of *MtYSL7* start codon were amplified using the primers indicated in Suppl. Table 1, then cloned in pDONR207 (Invitrogen) and transferred to pGWB3 (Nakagawa et al., 2007) using Gateway Cloning technology (Invitrogen). This led to the fusion of the promoter region of *MtYSL7* with the *gus* gene in pGWB3. An *A. rhizogenes* ARqua1 derived strain containing pGWB3::*MtYSL7*_*prom*_ vector was used for root transformation of *M. truncatula*. Transformed plants were transferred to sterilized perlite pots and inoculated with *S. meliloti* 2011. GUS activity was determined in 28 dpi plants as described (Vernoud et al., 1999).

### Immunohistochemistry and confocal microscopy

A DNA fragment of the full length *MtYSL7* genomic region and the 2 kb upstream of its start codon was amplified using the primers indicated in Suppl. Table 1 and cloned into the plasmid pGWB13 (Nakagawa et al., 2007) using the Gateway technology (Invitrogen). This plasmid fuses three C-terminal hemagglutinin (HA) epitopes in-frame. Hairy-root transformation was performed as previously described (Vernoud *et al*., 1999). Transformed plants were transferred to sterilized perlite pots and inoculated with *S. meliloti* 2011 containing a pHC60 plasmid that constitutively expresses GFP. Roots and nodules collected from 28 dpi plants were fixed by overnight incubation in 4 % paraformaldehyde, 2.5 % sucrose in PBS at 4 °C. After washing in PBS, nodules were included in 6 % agarose and cut in 100 μm sections with a Vibratome 1000 plus (Vibratome). Sections were dehydrated using methanol series (30, 50, 70, 100 % in PBS) for 5 min and then rehydrated. Cell walls were permeabilized with 4 % cellulase in PBS for 1 h at room temperature and with 0.1 % Tween 20 in PBS for 15 min. Sections were blocked with 5 % bovine serum albumin (BSA) in PBS before their incubation with an anti-HA mouse monoclonal antibody (Sigma) for 2 hours at room temperature. After washing, an Alexa594-conjugated anti-mouse rabbit monoclonal antibody (Sigma) was added to the sections for 1 h at room temperature. DNA was stained with DAPI after washing. Images were acquired with a confocal laser-scanning microscope (Leica SP8) using excitation lights at 488 nm for GFP and at 561 nm for Alexa 594.

### Acetylene reduction assay

Nitrogenase activity was measured by the acetylene reduction assay (Hardy et al., 1968). Nitrogen fixation was assayed in mutants and control plants at 28 dpi in 30 ml vials fitted with rubber stoppers. Each vial contained four or five pooled transformed plants. Three ml of air inside of the vial was replaced with 3 ml of acetylene. Tubes were incubated at room temperature for 30 min. Gas samples (0.5 ml) were analyzed in a Shimadzu GC-8A gas chromatograph fitted with a Porapak N column. The amount of ethylene produced was determined by measuring the height of the ethylene peak relative to background. Each point consists of two vials each. After measurements, nodules were recovered from roots to measure their weight.

### Metal content determination

Iron content was determined in shoots, roots, and nodules 28 dpi. Plant tissues were weighted and mineralized in 15.6 M HNO_3_ (trace metal grade) for 1 h at 80 °C and overnight at 20 °C. Digestions were completed with 2 M H_2_O_2_. Samples were diluted in 300 mM HNO_3_ prior to measurements. Element analyses were performed with Atomic Absorption Spectroscopy (AAS) in an AAnalyst 800 (Perkin Elmer), equipped with a graphite furnace. All samples were measured in duplicate.

### Bioinformatics

To identify *M. truncatula* YSL family members, BLASTN and BLASTX searches were carried out in the *M. truncatula* Genome Project site (http://www.jcvi.org/medicago/index.php). Protein sequences for tree construction were obtained from Phytozome (https://phytozome.jgi.doe.gov/pz/portal.html), Uniprot (http://www.uniprot.org/blast) and NCBI (https://blast.ncbi.nlm.nih.gov/Blast.cgi?PAGE=Proteins): *M. truncatula* MtYSL1 (Medtr1g077840); MtYSL2 (Medtr1g007540); MtYSL3 (Medtr3g092090); MtYSL4 (Medtr1g007580); MtYSL5 (Medtr6g077870); MtYSL6 (Medtr7g028250); MtYSL7 (Medtr3g063490), MtYSL8 (Medtr5g091600) y MtYSL9 (Medtr3g063520); *Arabidopsis thaliana* AtYSL1 (At4g24120), AtYSL2 (At5g24380), AtYSL3 (At5g53550), AtYSL4 (At5g41000), AtYSL5 (At3g17650), AtYSL6 (At3g27020), AtYSL7 (At1g65730), AtYSL8 (At1g48370); *Oryza sativa* OsYSL1 (Os01g0238700), OsYSL3 (Os05g0251900), OsYSL4 (Os05g0252000); OsYSL5 (Os04g0390600), OsYSL6 (Os04g0390500); OsYSL7 (Os02g0116300), OsYSL8 (Os02g0116400), OsYSL10 (Os04g0674600), OsYSL12 (Os04g0524600), OsYSL13 (Os04g0524500), OsYSL14 (Os02g0633300); OsYSL15 (Os02g0650300), OsYSL16 (Os04g0542800); OsYSL17 (Os08g0280300), OsYSL18 (Os01g0829900); *Zea mays* ZmYS1 (Zm00001d017429), ZmYSL2 (Zm00001d025977), ZmYSL6 (Zm00001d003941), ZmYSL11 (Zm00001d025888), ZmYSL12 (Zm00001d025887), ZmYSL14A (Zm00001d051193), ZmYSL17 (Zm00001d054042); *Brachypodium distachyon* BdYS1A (BRADI_3g50267), BdYS1B (BRADI_3g50263), BdYSL2 (BRADI_3g50260), BdYSL3 (BRADI_5g17230), BdYSL9 (BRADI_5g17210); BdYSL10 (BRADI_2g5395), BdYSL11 (BRADI_5g16190), BdSYL12 (BRADI_5g16170), BdYSL13 (BRADI_5g16160) and *Glycine max* GmYSL7 (GLYMA_11G203400), GLYMA_09G164500; GLYMA_16G212900.

Trees were constructed from a ClustalW multiple alignment of the sequences (http://www.ebi.ac.uk/Tools/msa/clustalw2), then analyzed by MEGA7 (Kumar et al., 2016) using a Neighbour-Joining algorithm with bootstrapping (1,000 iterations). Unrooted trees were visualized with FigTree (http://tree.bio.ed.ac.uk/software/figtree).

### Statistical tests

Data were analyzed with Student’s unpaired *t* test to calculate statistical significance of observed differences. Test results with *p*-values lower than 0.05 were considered as statistically significant.

## ACKNOWLEDGEMENTS

We would also like to acknowledge the other members of laboratory 281 at Centro de Biotecnología y Genómica de Plantas (UPM-INIA) for their support and feedback in preparing this manuscript.

