## Supplementary material for "*Medicago truncatula Yellow Stripe-Like7* encodes a peptide transporter required for symbiotic nitrogen fixation": Supp. Materials and Methods

### SUPPLEMENTAL MATERIAL AND METHODS

**Yeast complementation assays**

For metal complementation assays, *MtYSL7* coding sequence was amplified from cDNA obtained from RNA isolated from 28 dpi *M. truncatula* nodules using the primers listed in Supp. Table 1. Amplicon was cloned in pDONR207 (Invitrogen) and transferred to the destination vector pYES6CT (Invitrogen), using the Gateway Cloning technology (Invitrogen). Empty pYES6CT or pYES6CT::*MtYSL7* were co-transformed with vector pGEV-Trp to drive the expression of the Gal promoter of pYES6CT using β-estradiol (Gao and Pinkham, 2000). Transformations of *Saccharomyces cerevisiae* cells were performed by the lithium acetate method (Schiestl and Gietz, 1989), and selected in SD media by their autotrophy. Iron uptake tests were carried out using the strains DEY1451 (*MATa/MATa ade2/+ canl/canl his3/his3 leu2/leu2 trpl/trpl ura3/ura3*; (Dix et al., 1994)), and mutant in iron transport DEY1453 (*ade2/+ canl/canl his3/his3 leu2/leu2 trpl/trpl ura3/ura3 fet3-2::HIS3/fet3-2::HIS3 fet4-1::LEU2/fet4-1::LEU2*; (Eide et al., 1996)). For DEY1453 strain to grow without the two mutant transporters, the medium has to contain 25 mM sodium citrate (pH 4.2) and 50 μM $FeCl_3$. For complementation assays with iron and iron-nicotianamine, plates were made as described by (Chu et al., 2010). To test whether MtYSL7 was able to restore copper uptake, the parental strain BY4741 (MATa *his3 leu2 met15 ura3*; Yeast Knockout Collection, GE Dharmacon), and mutant in CTR1 copper transporter DY4741*Δctr1* (MATa *his3 leu2 met15 ura3 ctr1::ura3)* were used. For these complementation assays with copper or copper-nicotianamine, glycerol was used as a carbon source in place of dextrose and plates were made following the concentrations indicated by (DiDonato et al., 2004). Finally, the strains DY1457 (*MATa ade6 can1 his3 leu2 lys2 trp1 ura3*) and the mutant in two zinc transporters ZHY3 (*MATa ade6 can1 his3 leu2 lys2 trp1 ura3 zrt1::LEU2 zrt2::HIS3*; (Zhao and Eide, 1996, 1996)) were used to test zinc uptake. For complementation assays with zinc or zinc-nicotianamine, SD-Trp medium was made with zinc-free YNB (yeast nitrogen base) and with 1.4 μM $ZnSO_4$ containing or lacking 10 μM nicotianamine.

To analyse MtYSL7 IRON-MAN peptide uptake capabilities (Grillet et al., 2018), pDR196 and pDR196::*MtYSL7* were transformed in yeast strains YJL212C (OPT1) (Euroscarf) (BY4741; *MATa; ura3Δ0; leu2Δ0; his3Δ1; met15Δ0; YJL212c::kanMX4*) and BY4741 (Winston et al., 1995) were used. Growth experiments were performed using

SD media without nitrogen (N) using the peptides ENGGDDDDSGYDYAPAA, GDDDD, or APAA, as nitrogen source. SD media supplemented with 5 g/l $(NH_4)_2SO_4$ was used as a growth control media.

For glutathione transport assays, the parental line ABC733 (BY4741) (*MATα, his3 leu2 met15 ura3*) (Brachmann et al., 1998) and the mutant in glutathione transport ABC817 (*hgt1)* (*MATα, his3 leu2 met15 ura3 hgt1::LEU2*) (Bourbouloux et al., 2000) were used. The first of them was transformed with empty pDR196 and the mutant was transformed with empty pDR196 and pDR196::*MtYSL7.* Serial dilutions of each transformant were growth 2-3 days at 30°C in SD media where the unique source of sulfur was 100µM of glutathione (GSH). SD medium with all required amino acids and SD without sulfur and supplemented with methionine were used as growth control media.

**Iron localization assays**

Roots and nodules from 28 dpi plants were collected, fixed, and included in LR-white resin as previously described (Rodríguez-Haas et al., 2013). Thin sections from these nodules were obtained at Centro Nacional de Microscopía Electrónica, and the Perl-DAB method was used to visualize iron distribution (Roschzttardtz et al., 2009). Direct observation of sections was performed under a Zeiss Axiophot photomicroscope (Carl Zeiss, Oberkochen, Germany) with an attached digital camera (Leica DFC 420C, Heerburgg, Switzerland). A minimum of three nodules and roots per treatment and three sections per sample were examined.

**Bourbouloux A, Shahi P, Chakladar A, Delrot S, Bachhawat AK** (2000) Hgt1p, a high affinity glutathione transporter from the yeast *Saccharomyces cerevisiae*. J. Biol. Chem. **275:** 13259-13265

**Brachmann CB, Davies A, Cost GJ, Caputo E, Li J, Hieter P, Boeke JD** (1998) Designer deletion strains derived from *Saccharomyces cerevisiae* S288C: a useful set of

strains and plasmids for PCR-mediated gene disruption and other applications. Yeast **14:** 115-132

**Chu HH, Chiecko J, Punshon T, Lanzirotti A, Lahner B, Salt DE, Walker EL** (2010) Successful reproduction requires the function of *Arabidopsis* Yellow Stripe-Like1 and Yellow Stripe-Like3 metal-nicotianamine transporters in both vegetative and reproductive structures. Plant Physiol. **154:** 197-210

**DiDonato RJ, Jr., Roberts LA, Sanderson T, Eisley RB, Walker EL** (2004) *Arabidopsis* Yellow Stripe-Like2 (YSL2): a metal-regulated gene encoding a plasma membrane transporter of nicotianamine-metal complexes. Plant J. **39:** 403-414

**Dix DR, Bridgham JT, Broderius MA, Byersdorfer CA, Eide DJ** (1994) The FET4 gene encodes the low affinity Fe(II) transport protein of *Saccharomyces cerevisiae*. J. Biol. Chem. **269:** 26092-26099

**Eide D, Broderius M, Fett J, Guerinot ML** (1996) A novel iron-regulated metal transporter from plants identified by functional expression in yeast. Proc. Natl. Acad. Sci. U S A **93:** 5624-5628

**Gao CY, Pinkham JL** (2000) Tightly regulated, β-estradiol dose-dependent expression system for yeast. BioTechniques **29:** 1226-1231

**Grillet L, Lan P, Li W, Mokkapati G, Schmidt W** (2018) IRON MAN is a ubiquitous family of peptides that control iron transport in plants. Nature Plants **4:** 953-963

**Rodríguez-Haas B, Finney L, Vogt S, González-Melendi P, Imperial J, González-Guerrero M** (2013) Iron distribution through the developmental stages of *Medicago truncatula* nodules. Metallomics **5:** 1247-1253

**Roschzttardtz H, Conéjéro G, Curie C, Mari S** (2009) Identification of the endodermal vacuole as the iron storage compartment in the Arabidopsis embryo. Plant Physiol. **151:** 1329-1338

**Schiestl RH, Gietz RD** (1989) High efficiency transformation of intact yeast cells using single stranded nucleic acids as carrier. Curr. Genet. **16:** 339-346

**Winston F, Dollard C, Ricupero-Hovasse SL** (1995) Construction of a set of convenient Saccharomyces cerevisiae strains that are isogenic to S288C. Yeast **11:** 53-55

**Zhao H, Eide D** (1996) The yeast ZRT1 gene encodes the zinc transporter protein of a high-affinity uptake system induced by zinc limitation. Proc. Natl. Acad. Sci. U S A **93:** 2454-2458

**Zhao H, Eide D** (1996) The ZRT2 gene encodes the low affinity zinc transporter in *Saccharomyces cerevisiae*. J. Biol. Chem. **271:** 23203-23210
