## Supplementary material for "*Medicago truncatula Yellow Stripe-Like7* encodes a peptide transporter required for symbiotic nitrogen fixation": Supp. Figures

**SUPPLEMENTAL FIGURE S1**

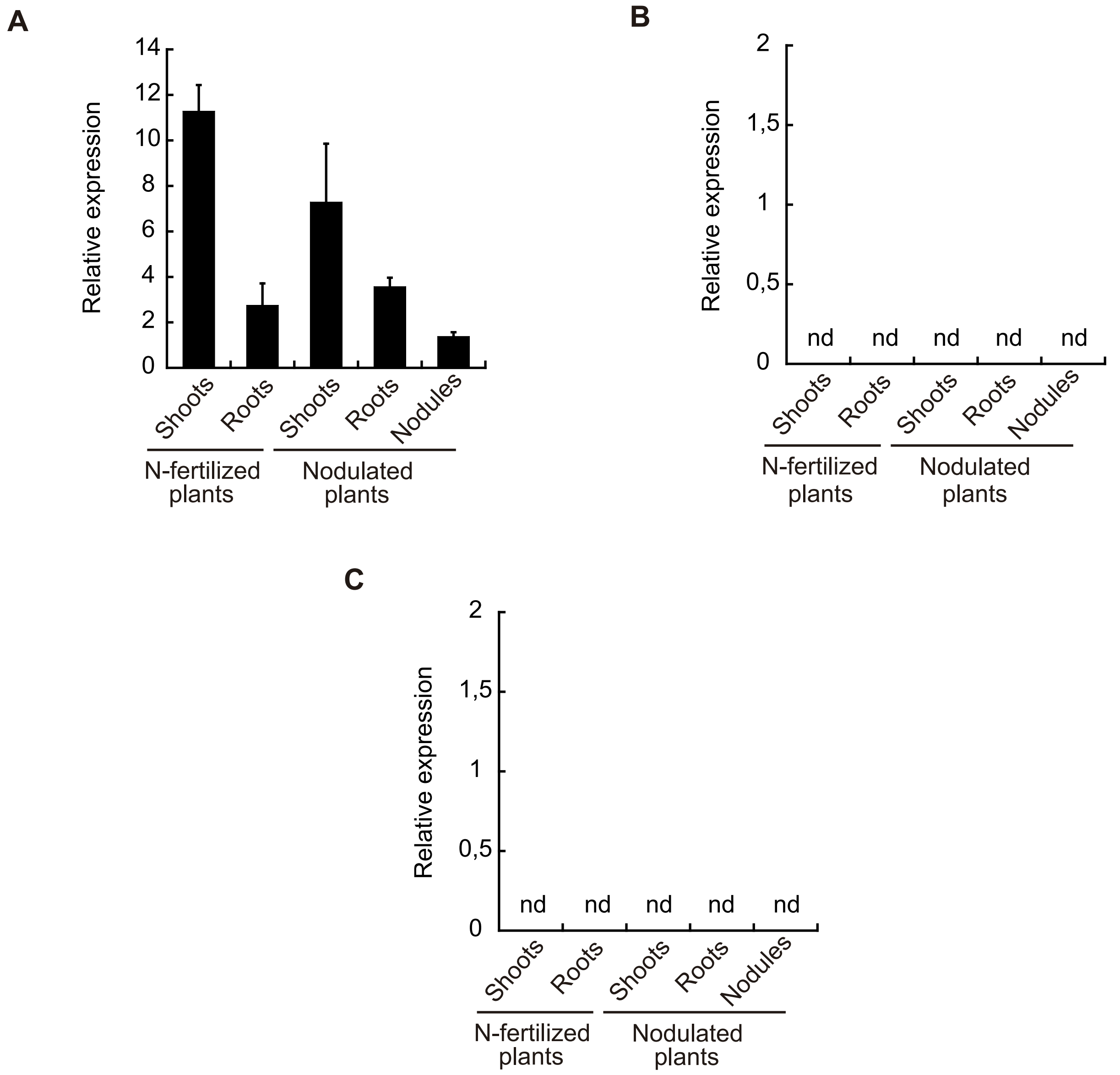


**Fig. S1.** Expression of *M. truncatula* Group III YSLs. A) *MtYSL5* expression relative to internal standard gene *ubiquitin carboxyl-terminal hydrolase.* Data are the mean ± SE of five independent experiments. B) *MtYSL8* expression relative to internal standard gene *ubiquitin carboxyl-terminal hydrolase*. Data are the mean ± SE of five independent expe-riments. C) *MtYSL9* expression relative to internal standard gene *ubiquitin carboxyl-terminal hydrolase.* Data are the mean ± SE of five independent experiments. n.d.: not detected.

### SUPPLEMENTAL FIGURE S2

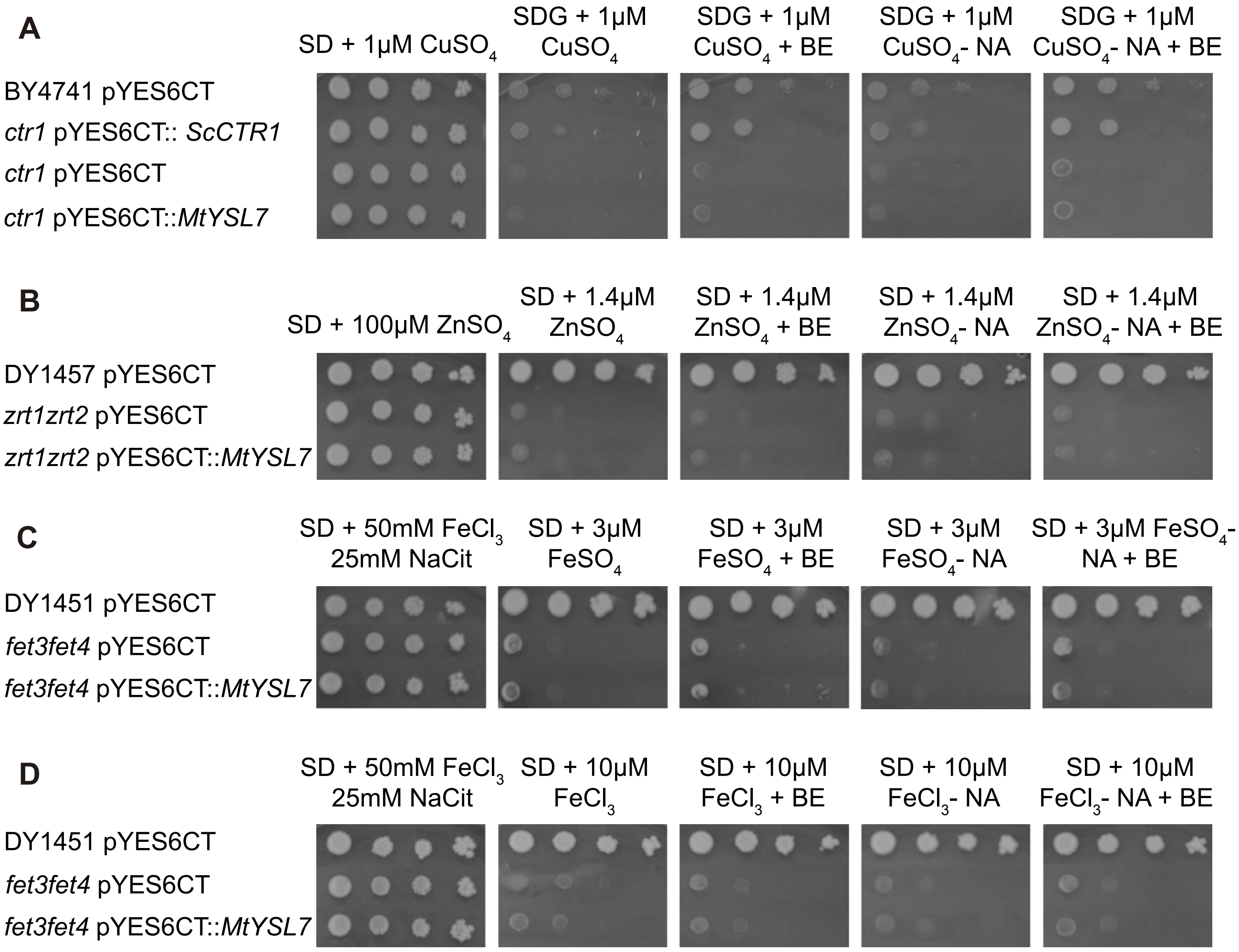


**Fig. S2.** MtYSL7 metal nicotianamine transport assays. A) Copper or copper-nicotianamine (NA) uptake assay. Wild type *S. cerevisiae* strain BY4741 was transformed empty pYES6CT vector, while the *ctr1* copper uptake mutant was transformed with the empty pYES6CT, pYES6CT containing *ScCTR1*, or pYES6CT containing MtYSL7 cDNA. All of these strains were co-transformed with pGEV vector. Serial dilution were placed in SD or SDG (SD-glycerol) media supplemented with the indicated molecules. *Gal* promoter was induced with the addition of 10 nM of β-estradiol (BE). B) Zinc or zinc-nicotianamine (NA) uptake assay. Wild type *S. cerevisiae* strain DY14571 was transformed empty pYES6CT vector, while the *zrt1/zrt2* zinc uptake double mutant was transformed with the empty pYES6CT, or pYES6CT containing *MtYSL7* cDNA. All of these strains were co-transformed with pGEV vector. Serial dilution were placed in SD media supplemented with the indicated molecules. *Gal* promoter was induced with the addition of 10 nM of β-estradiol (BE). C) Iron(II) or $Fe^{2+}$-nicotianamine (NA) uptake assay. Wild type *S. cerevisiae* strain DY1451 was transformed empty pYES6CT vector, while the *fet3/fet4* iron uptake double mutant was transformed with the empty pYES6CT, or pYES6CT containing *MtYSL7* cDNA. All of these strains were co-transformed with pGEV vector. Serial dilution were placed in SD media supplemented with the indicated molecules. *Gal* promoter was induced with the addition of 10 nM of β-estradiol (BE). 25 mM sodium citrate (NaCit) was added to the medium to allow growth of *fet3/fet4* mutant strains. D) Iron(III) or $Fe^{3+}$-nicotianamine (NA) uptake assay. Wild type *S. cerevisiae* strain DY1451 was transformed empty pYES6CT vector, while the fet3/fet4 iron uptake double mutant was transformed with the empty pYES6CT, or pYES6CT containing *MtYSL7* cDNA. All of these strains were co-transformed with pGEV vector. Serial dilution were placed in SD media supplemented with the indicated molecules. *Gal* promoter was induced with the addition of 10 nM of β-estradiol (BE). 25 mM sodium citrate (NaCit) was added to the medium to allow growth of fet3/fet4 mutant strains.

### SUPPLEMENTAL FIGURE S3

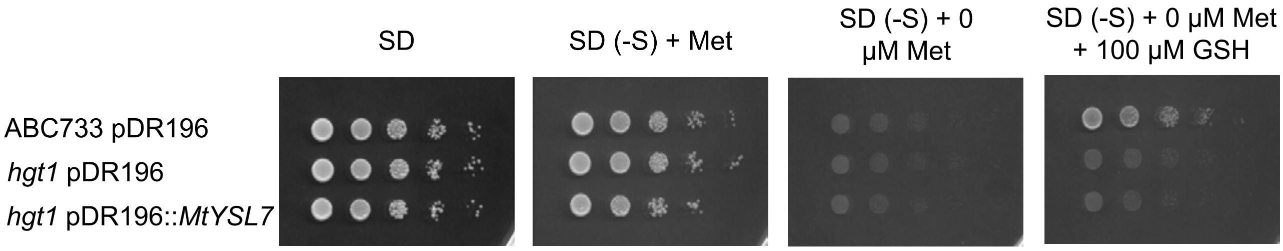


**Fig. S3.** MtYSL7 glutathione transport assays. *S. cerevisiae* strain ABC733 was transformed with empty pDR196 vector, while *hgt1* mutant was transformed with either the empty pDR196 or pDR196 expressing *MtYSL7* cDNA. Serial dilutions (10x) of each transformant were placed in SD medium, SD medium using either methionine (Met) or glutathione (GSH) as sole sulfur source.

**SUPPLEMENTAL FIGURE S4**

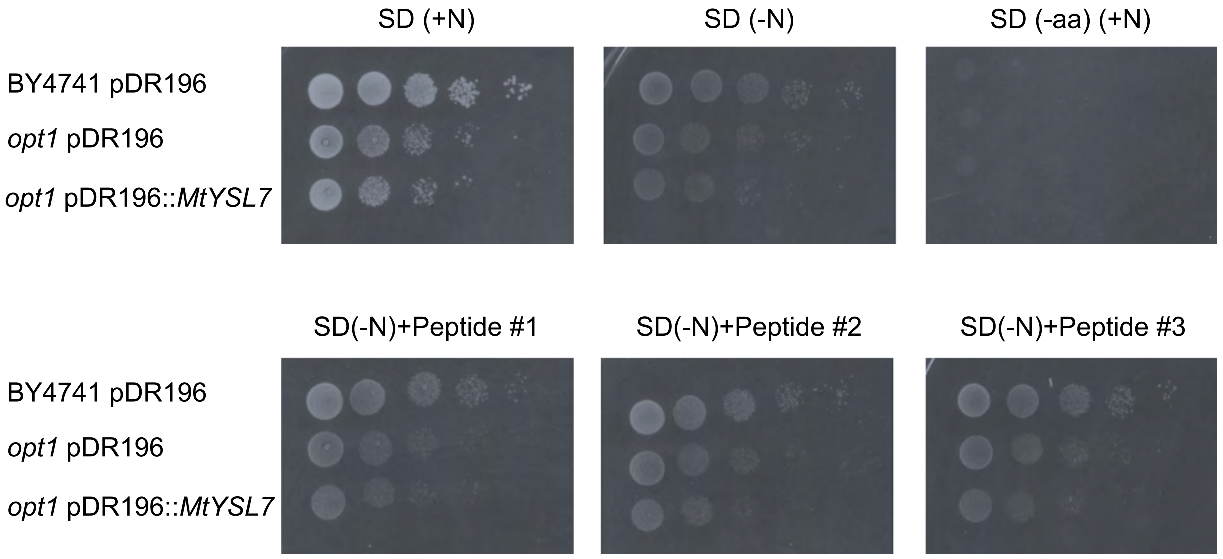


**Fig. S4.** MtYSL7 IMA peptides transport assays. Parental yeast strain BY4741 was transformed with empty pDR196 vector, while the strain *opt1*, mutant in the oligopeptide transporter ScOPT1, was transformed with either the empty pDR196 vector, or with pDR196 containing the coding sequence of *MtYSL7*. Serial dilutions (10x) were grown on SD media supplemented with one of the following peptides as sole nitrogen source: ENGGDDDDSGYDYAPAA (Peptide 1), GDDDD (Peptide 2), or APAA (Peptide 3).

**SUPPLEMENTAL FIGURE S5**

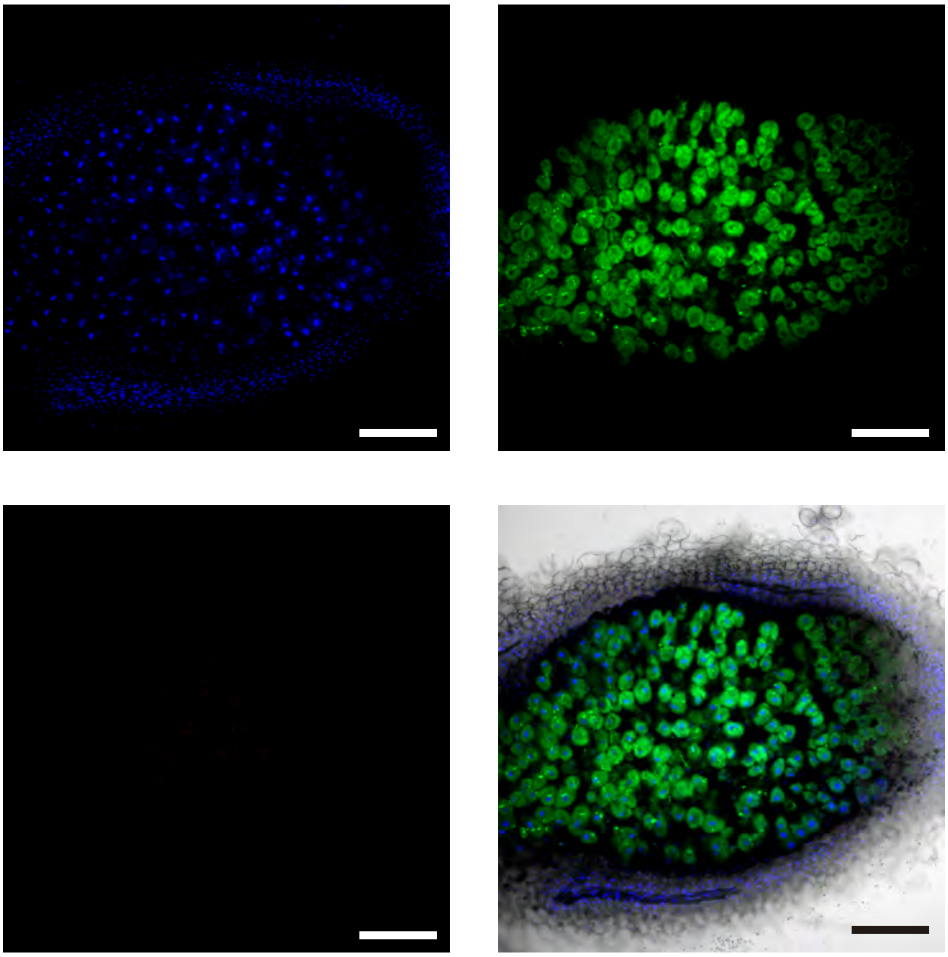

**Fig. S5.** Autofluorescence control for Alexa594. Longitudinal section of a 28 dpi *M. truncatula* nodule expressing *MtYSL7-HA* under its own promoter. No primary anti-HA antibody was used, but samples were incubated with secondary Alexa594-conjugated antibody and DNA stained with DAPI. Transformed plants were inoculated with a GFP-expressing *S. meliloti* strain. Top left panel corresponds to DAPI signal (blue), top right to the GFP emission (green), and lower left to the Alexa 594 emission with the same settings used in all confocal imaging. These three channels are overlaid with the bright field image in the lower right panel. Bars = 200 μm.

**SUPPLEMENTAL FIGURE S6**

**A**

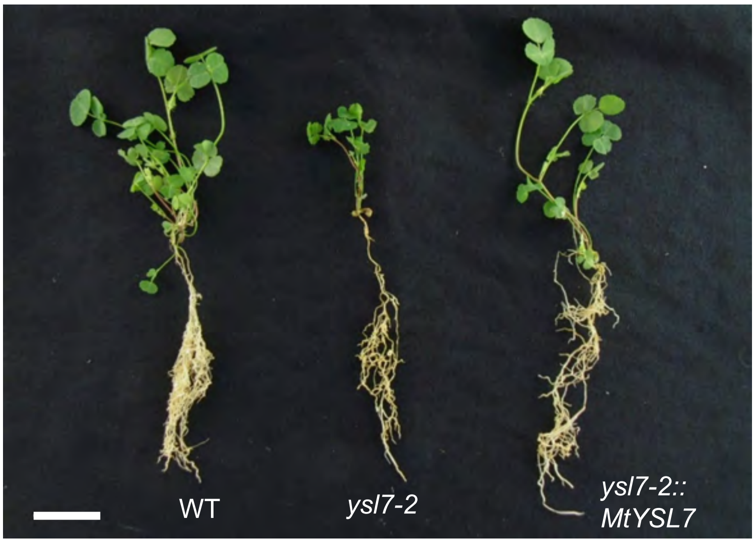


**B**

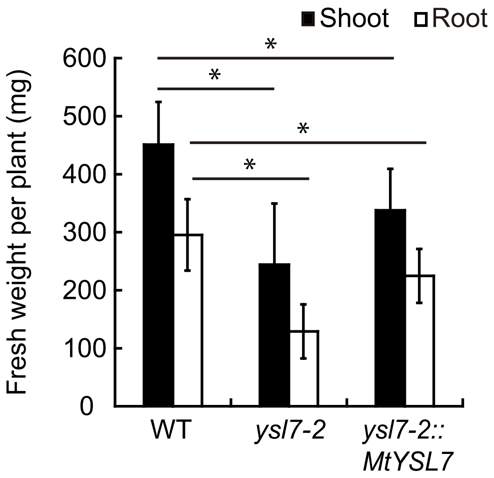


**Fig. S6.** Genetic complementation of the *ysl7-2* phenotype under non-symbiotic conditions. A). Growth of representative wild type, *ysl7-2*, and *ysl7-2* transformed with a wild type copy of *MtYSL7* regulated by its own promoter (*ysl7-2*::*MtYSL7*) plants. Bar = 3 cm. B) Fresh weight of shoots and roots of wild type, *ysl7-2*, and *ysl7-2::MtYSL7* plants. Data are the mean ± SE (n = 6 plants). * indicates statistical significance ($p < 0.05$).

**SUPPLEMENTAL FIGURE S7**

**A**

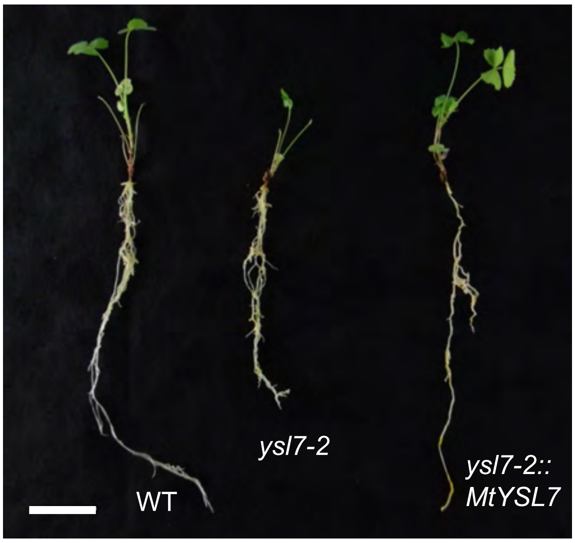


**B**

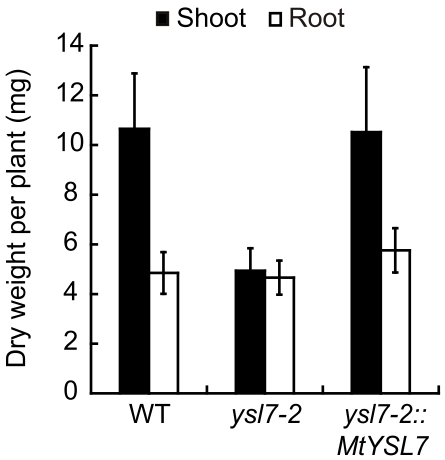


**C**

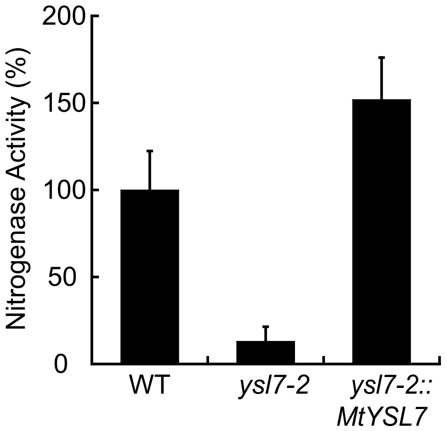


**Fig. S7.** Genetic complementation of the *ysl7-2* phenotype under symbiotic conditions. A) Growth of representative wild type, *ysl7-2*, and *ysl7-2* transformed with a wild type copy of *MtYSL7* regulated by its own promoter (*ysl7-2::MtYSL7*) plants. Bar = 3 cm B) Dry weight of shoots and roots of 28 dpi plants. Data are the mean ± SE (n = 5 plants). C) Nitrogenase activity in 28 dpi nodules from wild type, *ysl7-2* and *ysl7-2:: MtYSL7* plants. Data are the mean ± SE measured in duplicate from two sets of five pooled plants. 100 % = 0.115 nmol ethylene h-1 plant-1. * indicates statistical significance ($p < 0.05$).

**SUPPLEMENTAL FIGURE S8**

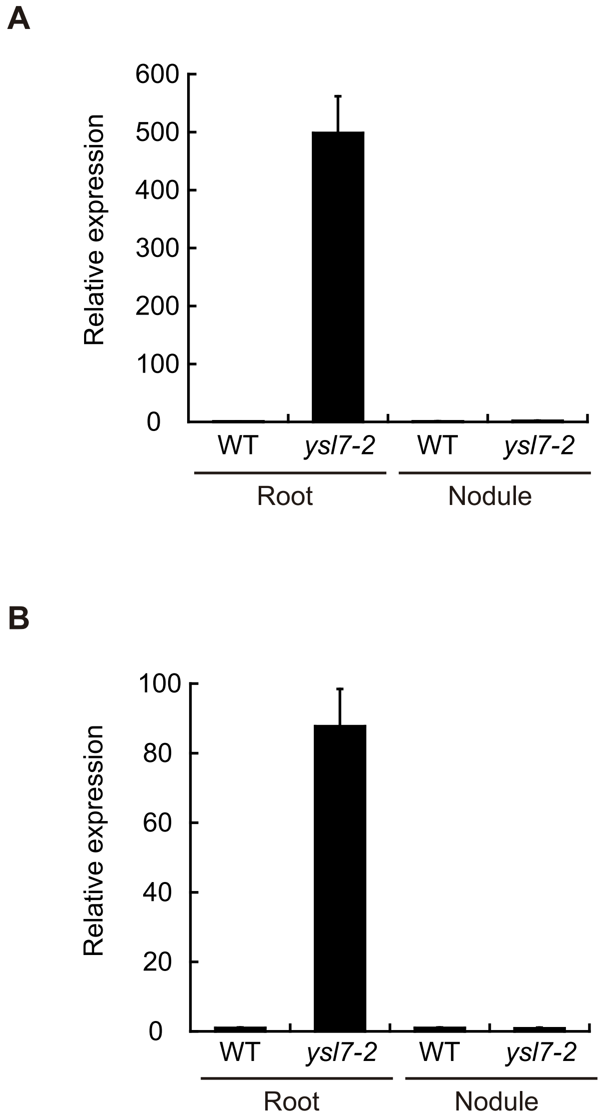


**Fig. S8.** Expression of *M. truncatula* iron homeostasis genes A) *MtFRO1* expression relative to internal standard gene *ubiquitin carboxyl-terminal hydrolase* in 28 dpi roots and nodules from wild type (WT) and *ysl7-2* plants. Data bars are standardized to wild-type values (1) and show the mean ± SE of two independent experiments. B) *MtNramp1* expression relative to internal standard gene *ubiquitin carboxyl-terminal hydrolase* in 28 dpi roots and nodules from wild type (WT) and *ysl7-2* plants. Data bars are standardized to wild-type values (1) and show the mean ± SE of two independent experiments.

**SUPPLEMENTAL FIGURE S9**

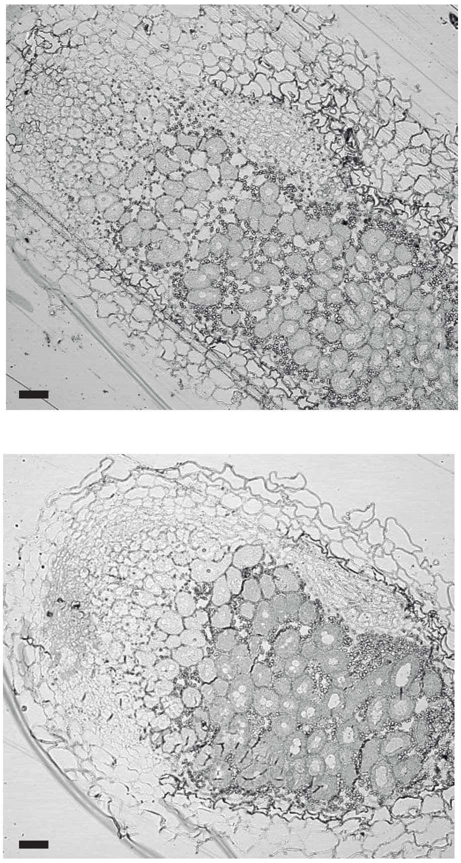

**Fig. S9.** Iron distribution in 28 dpi wild type (top panel) and ysl7-2 (bottom panel) nodules. Bars = 50 µm.

### SUPPLEMENTAL FIGURE S10

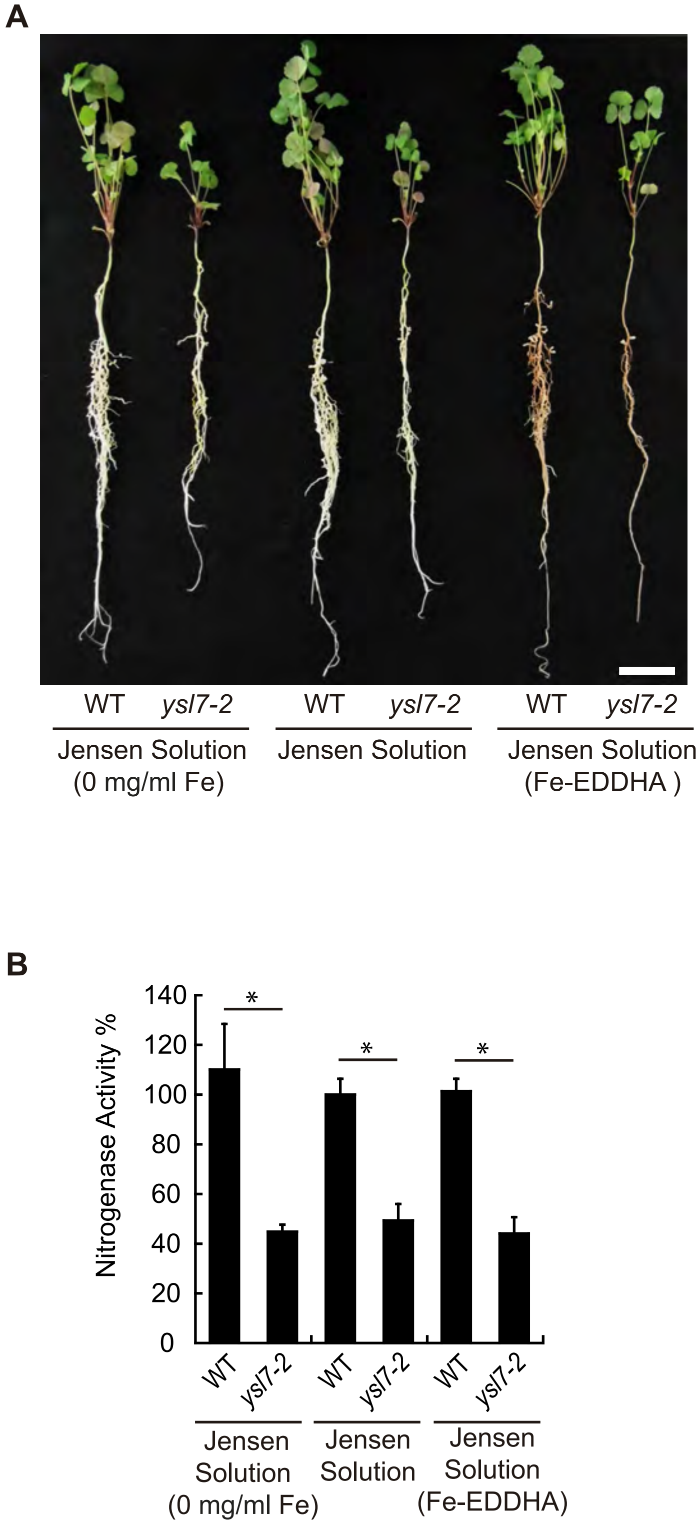


**Fig. S10.** Effect of changing iron concentrations on the *ysl7-2* phenotype. A) Growth of representative wild type and *ysl7-2* plants watered with standard nutrient solution (Jensen solution), with a modified nutrient solution without adding any iron (0 mg/ml Fe), and one fortified with 0.5 g/l of Fe-EDDHA. Bar = 3 cm. B) Nitrogenase activity in 28 dpi nodules from wild type or *ysl7-2* plants watered with standard nutrient solution (Jensen solution), with a modified nutrient solution without adding any iron (0 mg/ml Fe), and one fortified with 0.5 g/l of Fe-EDDHA. Data are the mean ± SE measured in duplicate from two sets of five pooled plants. 100 % = 0.645 nmol ethylene $h^{-1}$ $plant^{-1}$. * indicates statistical significance ($p < 0.05$).

### SUPPLEMENTAL FIGURE S11

A

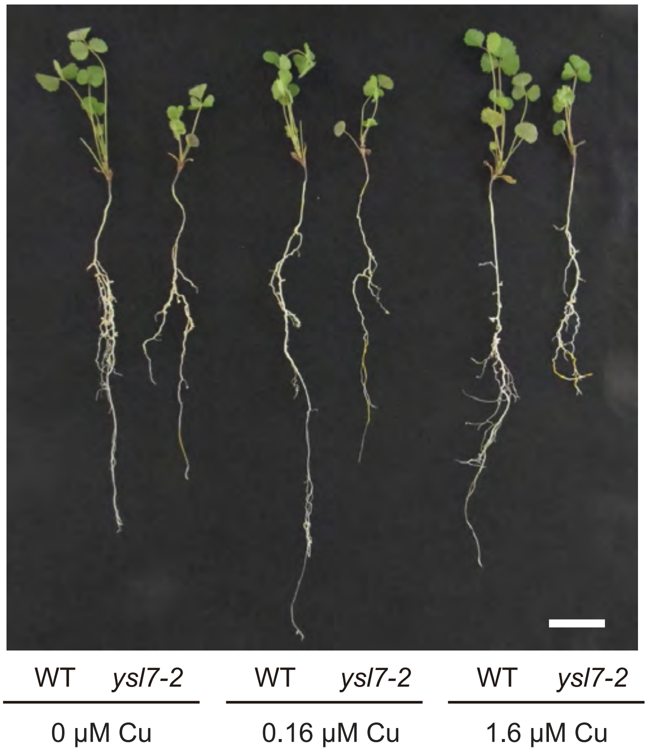


B

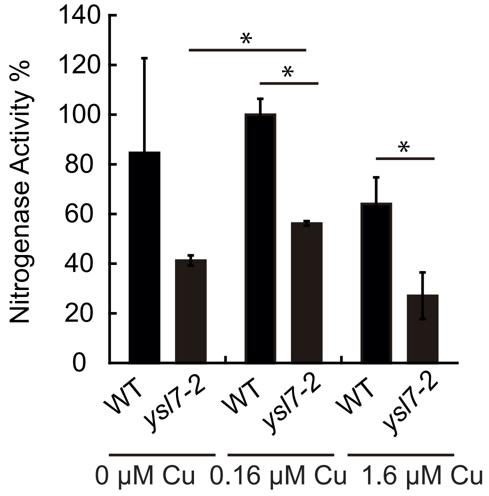


**Fig. S11.** Effect of changing copper concentrations on the *ysl7-2* phenotype. A) Growth of representative wild type and *ysl7-2* plants watered with standard nutrient solution (Jensen solution), with a modified nutrient solution without adding any copper (0 µM Cu), and one fortified with 1.6 µM Cu. Bar = 3 cm. B) Nitrogenase activity in 28 dpi nodules from wild type or *ysl7-2* plants watered with standard nutrient solution (Jensen solution), with a modified nutrient solution without adding any copper (0 µM Cu), and one fortified with 1.6 µM Cu. Data are the mean ± SE measured in duplicate from two sets of five pooled plants. 100 % = 0.456 nmol ethylene $h^{-1}$ $plant^{-1}$. * indicates statistical significance ($p < 0.05$).
