## Supplementary material for "*Medicago truncatula Yellow Stripe-Like7* encodes a peptide transporter required for symbiotic nitrogen fixation": Supp. Table

Supplemental Table 1. Primers used in this study

| Primer | Sequence | Use |
|---|---|---|
| MtUb v4qF | ATTCTTCACATGCGGCGATTTAC | qPCR of *ubiquitin carboxyl-terminal hydrolase* |
| MtUb v4qR | TTTCTCATTTGCTTTTGGTGTTGG | qPCR of *ubiquitin carboxyl-terminal hydrolase* |
| YSL7qF2 | CGTCGAAAGCCATGACAGTGGT | qRT-PCR of *MtYSL7* |
| YSL7qR2 | ACCGACAACATGAGACTCACGAA | qRT-PCR of *MtYSL7* |
| YSL8qF_v2 | CGTGACGTCGAAACCTCCGA | qRT-PCR of *MtYSL8* |
| YSL8qR_v2 | ATGATACTCACTACTATTGCCCTAACC | qRT-PCR of *MtYSL8* |
| YSL9qF | CAGTTGCGCCGGTGTTGAAA | qRT-PCR of *MtYSL9* |
| YSL9qR_v2 | TCACCGACAACATGAGACTTACA | qRT-PCR of *MtYSL9* |
| YSL5qF | GCTCTTTTTGGAGTTAAGCCTCCC | qRT-PCR of *MtYSL5* |
| YSL5qR | GGCAGATTTACAATGCATCAACAAGC | qRT-PCR of *MtYSL5* |
| 5MtNRAMP1-1533 | AAGGGGTTTGTAAACGATGGTCAGT | qRT-PCR of *MtNramp1* |
| 3MtNRAMP1-1659 | GCTCTCTAAAATGACTTGAAAGCAA | qRT-PCR of *MtNramp1* |
| 5MtFRO1qF | GGTGACACGTGGATCATCTG | qRT-PCR of *MtFRO1* |
| 3MtFRO1qR | TTGCAATCCACAGGAACAAA | qRT-PCR of *MtFRO1* |
| 5MtYSLpGW | GGGGACAAGTTTGTACAAAAAAGCAG GCTCTTAGGGAGGTTTTTTTAGAG | Cloning of *MtYSL7* promoter and full sequence in pGWB3/13 |
| 3MtYSLpGW | GGGGACCACTTTGTACAAGAAAGCTG GGTTATGTTTACAATGAAATTGA | Cloning of *MtYSL7* promoter in pGWB3 |
| 3MtYSLGW | GGGGACCACTTTGTACAAGAAAGCTG GGTTATGTTCTAAGAAGGCATCAAC | Cloning of *MtYSL7* promoter, full sequence and coding sequence for fused to 3xHA or GFP in C- terminous. |
| 5MtYSL7+1GW-C | GGGGACAAGTTTGTACAAAAAAGCAG GCTATGTCTTCTGAACCCCATCGA | Cloning of *MtYSL7* coding sequence in Gateway vectors to fuse them to GFP in C-terminous |
| 5MtYSL7+1GW-N | GGGGACAAGTTTGTACAAAAAAGCAG GCTTCATGTCTTCTGAACCCCATCGA | Cloning of *MtYSL7* coding sequence in Gateway vectors to fuse them to GFP in N-terminous |
| 3MtYSL7+3996GW-N | GGGGACCACTTTGTACAAGAAAGCTG GGTTCAATGTTCTAAGAAGGCATCAAC | Cloning of *MtYSL7* coding sequence in Gateway vectors to fuse them to GFP in N-terminous |
| 5MtYSL7+1_PstI | AAAACTGCAGATGTCTTCTGAACCCCA TCGA | Cloning of *MtYSL7* coding sequence in pDR196 |
| 3MtYSL7+4002_XhoI | AAAACTCGAGTCAATGTTCTAAGAAG GC | Cloning of *MtYSL7* coding sequence in pDR196 |
| YSL7.2-Fv.2 | CGAAAGCCATGACAGTGGT | Genotyping of Tnt1 mutant lines |
| YSL7.2-Rv.2 | AGGGGGTAATTAGATTAAACCAAA | Genotyping of Tnt1 mutant lines. |
